# Induced Pluripotent Stem Cell-derived CAR-Macrophage Cells with Antigen-dependent Anti-Cancer Cell Functions for Liquid and Solid Tumors

**DOI:** 10.1101/2020.03.28.011270

**Authors:** Li Zhang, Lin Tian, Xiaoyang Dai, Hua Yu, Jiajia Wang, Anhua Lei, Wei Zhao, Yuqing Zhu, Zhen Sun, Hao Zhang, George M. Church, He Huang, Qinjie Weng, Jin Zhang

## Abstract

The Chimera antigen receptor (CAR)-T cell therapy has gained great success in the clinic. However, there are still major challenges for its wider applications in a variety of cancer types including lack of effectiveness due to the highly complex tumor microenvironment, and the forbiddingly high cost due to personalized manufacturing procedures. In order to overcome these hurdles, numerous efforts have been spent focusing on optimizing Chimera Antigen Receptors, engineering and improving T cell capacity, exploiting features of subsets of T cell or NK cells, or making off-the-shelf universal T cells. Here, we developed induced pluripotent stem cells (iPSCs)-derived, CAR-expressing macrophage cells (CAR-iMac). These cells showed antigen-dependent macrophage functions such as expression and secretion of cytokines, polarization toward the pro-inflammatory/anti-tumor state, and phagocytosis of tumor cells, as well as some *in vivo* anti-cancer cell activity for both liquid and solid tumors. This technology platform for the first time provides an unlimited source of iPSC-derived engineered CAR-macrophage cells which could be utilized to eliminate cancer cells or modulate the tumor microenvironment in liquid and solid tumor immunotherapy.

**One sentence summary:** We developed CAR-expressing iPSC-induced macrophage cells that have antigen-dependent phagocytosis and pro-inflammatory functions and anti-cancer cell activity for both liquid and solid tumor cells.

## Introduction

Human iPSCs derived from patients’ own cells can be differentiated to multiple cell types and used for disease modeling, drug screening cell models and cell-based therapies^1^. Recently, engineered T cells have shown great success in treating cancers in the clinic^2^. However, there are a few outstanding challenges for engineering primary immune cells, including variable transduction efficiency, highly heterogeneous genetic outcomes upon editing the genome of the targeted cells, and limited cell resources for certain cases such as from hematological cancer patients for which immune cell itself is transformed. iPSC-derived immune cells, due to their flexibility of expansion and genome editing at the iPSC stage^3, 4^, in theory have advantages in dealing with those challenges above, and have been proved to be effective in treating B cell and ovarian cancer cells in pre-clinical settings such as iPSC differentiated T cells^5^ and NK cells^6^. Other than lymphoid lineage cytotoxic cells, a larger number of myeloid cells exist in the tumor microenvironment^7^. Recently, myeloid cells such as macrophages have been utilized as a type of effector cells to combat cancer cells by means of their phagocytosis function^8^. However, the immortalized cell lines used in the previous study are not applicable to clinical settings, and bone marrow-derived macrophages are not easily engineered, thus leaving iPSC-derived macrophage cells as a great source for myeloid cell-based cancer immunotherapy. In addition to phagocytosis function, it is believed that macrophages in a polarized pro-inflammatory state can alter the tumor microenvironment through secreting cytokines to stimulate T cells^9^. Moreover, it was reported that tumor macrophages transition to a pro-inflammatory polarized state upon immune-checkpoint therapy (ICT) treatment in animal models^10^, and an inflammatory signature of tumor samples predicts clinical benefits in ICT-treated patients^11^. Applying the above features of macrophages to combat tumor cells is currently under intensive investigation largely by targeting endogenous macrophages in *vivo*^12^, but not much effort has been spent on adoptive transfer therapies using monocyte/macrophages^13^, partly due to the difficulties in obtaining or expanding enough cells, and the low efficiency in engineering them in order to harness these cells for combating cancer. Here, we used human iPSCs to introduce a CD19 or mesothelin-targeting CAR, and differentiated them into macrophages, a product we named as CAR-iMac. We thoroughly characterized these cells with bulk/single cell RNA-sequencing and functional analyses, and showed they are heterogeneous and functional macrophage cells. Upon challenge with antigen-expressing cancer cells, CAR-iMac cells show antigen-dependent increase of phagocytosis, strong induction of antigen-dependent cytokine expression and secretion, and polarization toward a more inflammatory M1-like state. Transplanting CD19 or Mesothelin CAR-iMac cells to liquid and solid tumor models showed some *in vivo* anti-tumor effect. This CAR-iMac system provides a proof-of-principle platform to use pluripotent stem cells to obtain a large amount of engineered macrophage cells with antigen-dependent functions for adoptive transfer-based immune cell therapies.

## Results

### Engineering a CAR-expressing iPSC line for producing macrophages

We aim to develop engineered macrophages that have the following features: (1) the differentiated macrophages are capable of phagocytosing tumor cells and producing cytokines that may alter the tumor microenvironment; (2) these functions should be antigen-dependent; and (3) the engineered cells should have a high percentage of transgene expression and can be produced in large scale (Fig. 1a). First, we started from deriving iPSCs from Peripheral blood mononuclear cells (PBMC) of a healthy donor with non-integrable episomal vectors encoding OCT4, SOX2, KLF4, L-MYC and LIN28A (Supplementary Fig. 1a,b). Isolated single iPSC clones showed pluripotent markers OCT3/4 and LIN28A/B expression (Supplementary Fig. 1c), and formed teratomas when injected to immune-deficient mice (Supplementary Fig. 1d). We originally designed a CD19 CAR supposedly more suitable for macrophages by replacing the 4-1BB and CD3ζ intracellular domains (T-CAR) with the macrophage CD86 and FcγRI intracellular domains (M-CAR) (Supplementary Fig. 2a), and transduced M-CAR and T-CAR into the THP-1 human monocyte cell line with lentivirus and achieved comparable levels of expression (Supplementary Fig. 2b,c). To our surprise, even though the M-CAR-transduced cells showed increased level of cytokine expression and secretion when encountering the antigen CD19-expressing K562 cells compared with the untransduced control, the increase was more significant for the T-CAR-transduced cells (Supplementary Fig. 2d,e), suggesting T-CAR is capable of stimulating monocyte/macrophage intracellular signaling and activating macrophage specific genes. Based on these results, the CD19-targeting T-CAR transgene (named as CAR thereafter) was introduced into the iPSCs by lentiviral transduction (Fig. 1a and Supplementary Fig. 2f). qPCR showed mRNA of the CAR transgene was highly expressed in these iPSCs (Supplementary Fig. 2g). We thus engineered a CAR-expressing iPSC line amenable for producing differentiated products with antigen-dependent functions.

**Fig. 1.**
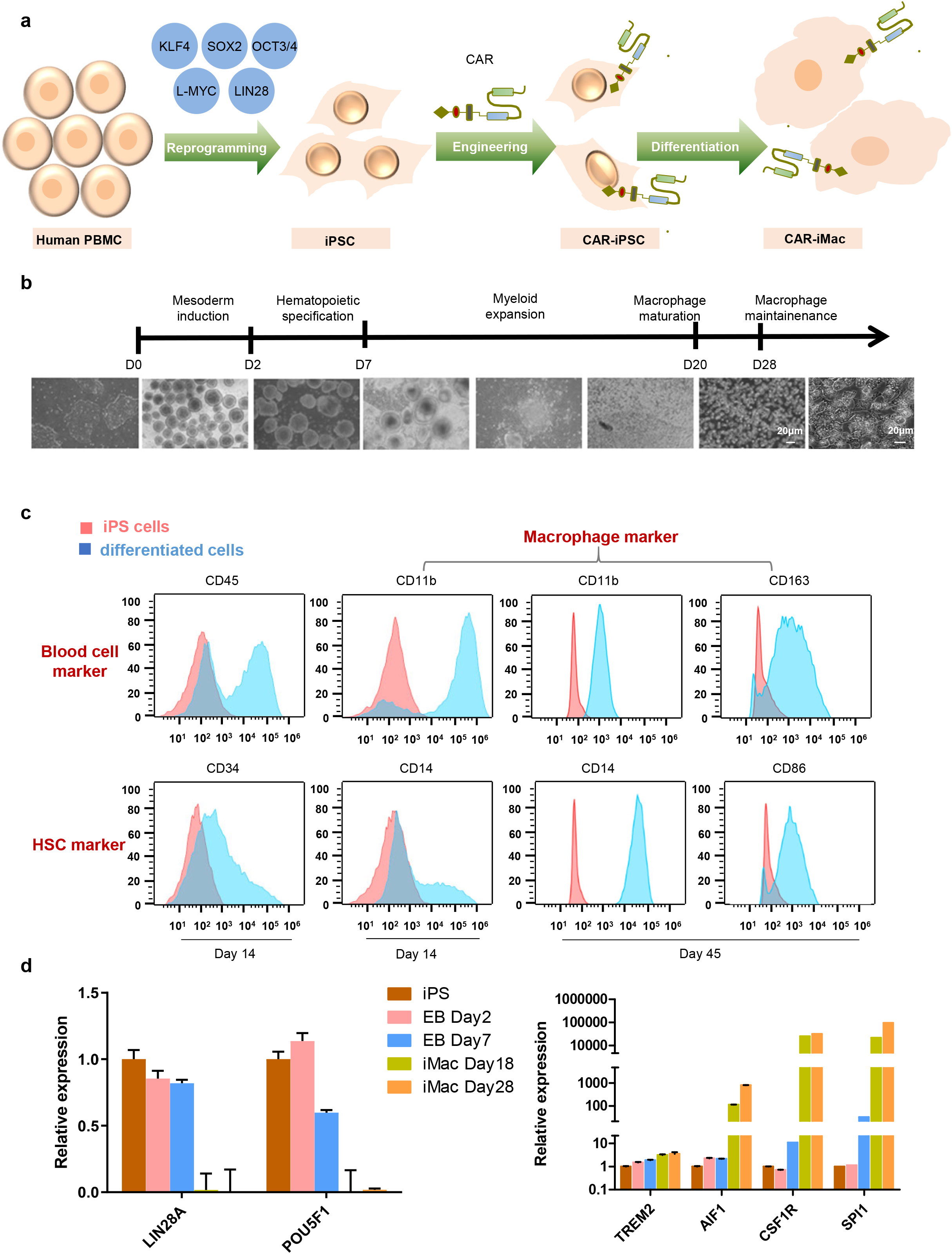
Differentiation of CAR-iPSCs into macrophages. **a**, Schematics showing the procedures of PBMC reprogramming, introduction of chimera antigen receptor and differentiation to macrophage cells. **b**, Overview of the differentiation protocol to derive macrophages from iPSCs. Representative microscopic pictures showing cells at different stages during differentiation. The last picture is a magnified filed for matured macrophages attached at the bottom of the plate at day 28. **c**, Flow cytometry analysis of iPSC-derived cells at different stages of differentiation with stage specific markers. **d**, qRT-PCR showing pluripotent marker gene and key macrophage marker gene expression at different stages of differentiation. n=3, error bar: standard error of the mean.

### CAR-expressing iPSCs can differentiate to macrophage cells

Next, we established a protocol of myeloid/macrophage differentiation to induce iPSCs toward myeloid cell lineages (Fig. 1b). After a period of mesoderm induction, hematopoietic specification, myeloid expansion, and macrophage maturation, differentiated cells showed typical macrophage marker gene expression such as CD14, CD11b, CD163 and CD86 (Fig. 1c), as well as CAR expression on cell surface detected with an antibody against CD19 CAR (Supplementary Fig. 2h). Consistently, key macrophage genes were induced such as *AIF1, CSF1R* and *SPI1,* whereas pluripotent marker genes disappeared such as *LIN28A* and *POU5F1* (Fig. 1d). RNA-sequencing using undifferentiated cells, early-day precursor cells and 18-day, 28-day 38-day differentiated cells showed that iPSCs clustered with precursor cells, and late-day differentiated cells clustered with primary macrophage cells, or iPSC-differentiated macrophage cells from previous studies^14, 15^ (Fig. 2a,b). However, it is challenging to classify iMac to M0, M1 or M2 types of macrophages, likely because of the heterogeneous feature of the primary and differentiated cells from the published bulk RNA-sequencing data. Similarly, selected M1 and M2 marker genes both showed high expression determined by qRT-PCR (Supplementary Fig. 3a, b). GO and KEGG pathway analyses showed strong enrichment of innate immunity-related functions in 18 or 28-day differentiated cells such as: “positive regulation of cytokine production”, “regulation of innate immune responses” and “regulation of phagocytosis” (Fig. 2c, d). Quantitative PCR also validated that the differentiated cells expressed high levels of Toll-like receptor, Fc receptor and cytokine receptor genes, HLA genes, NF-κB pathway genes, and costimulatory signaling genes, etc (Supplementary Fig. 3c-h), all of which are required for certain specific monocyte/macrophage/dendritic cell (DC) functions. Importantly, these genes are expressed in comparable levels with primary PBMC-derived macrophage cells (Supplementary Fig. 3i). Together, these data show CAR-expressing iPSCs can differentiate to myeloid lineage and macrophage-like cells.

**Fig. 2.**
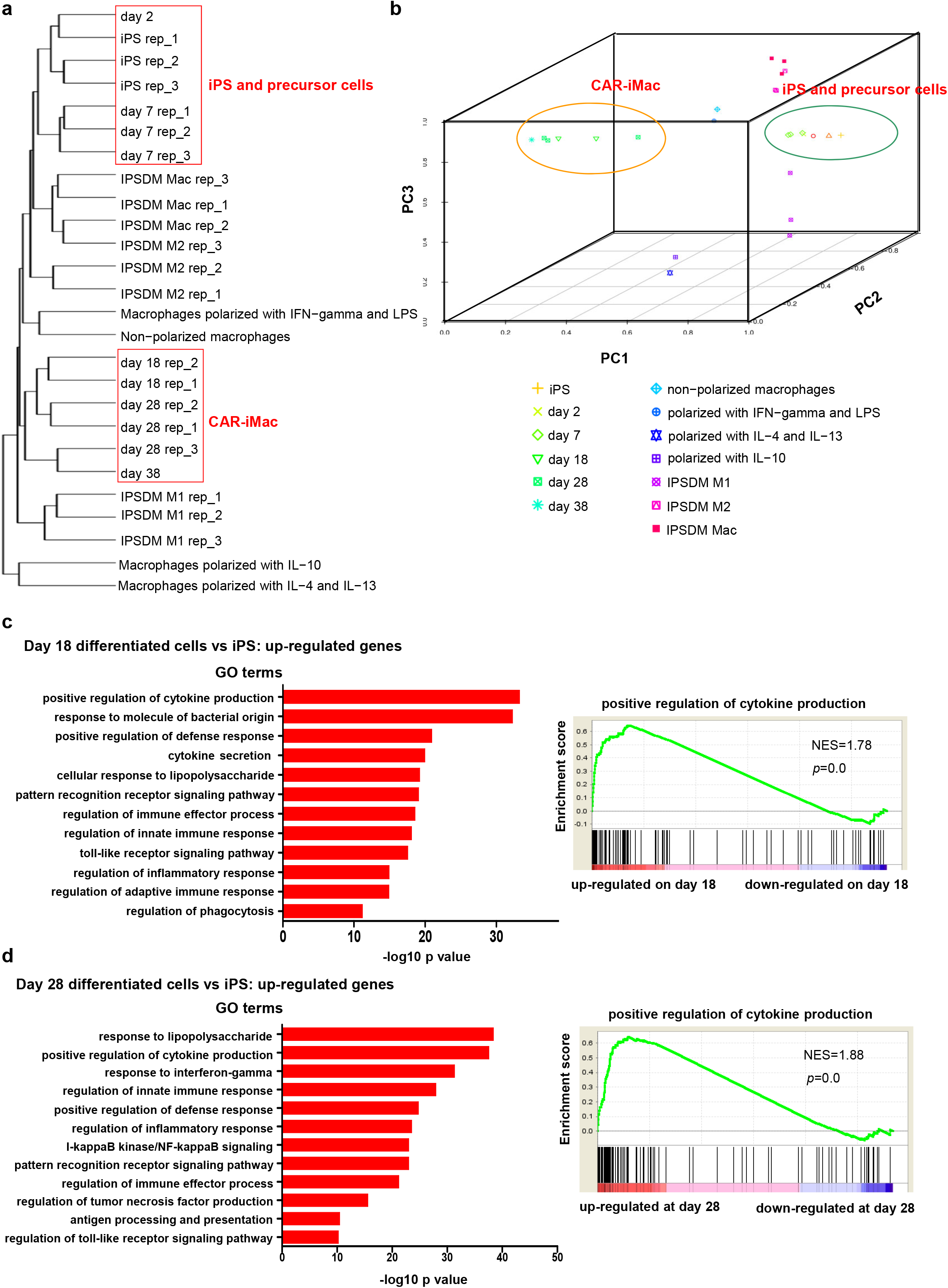
Transcriptional profiling of iPSC-derived CAR-iMac cells. **a**, Hierarchical clustering of transcriptomes of iPSCs, differentiated cells, and primary and stem cell-differentiated macrophages in different states. **b**, Principle component analysis (PCA) of the same samples as in **a**. **c**, Top GO terms enriched in genes up-regulated on day 18 differentiated cells compared with iPSCs. Right panel is an example of GSEA analysis of one GO term “Positive regulation of cytokine production”. NES: normalized enrichment score. P=0: p-value is a very small number. **d**, Top GO terms enriched in genes up-regulated on day 28 compared with iPSCs, and GSEA analysis of “Positive regulation of cytokine production”.

### Single cell RNA-seq analysis of CAR-iMac reveals a differentiation trajectory mainly toward M2 macrophage and dendritic cells

To dissect heterogeneity of these CAR-iMac cells, we performed single cell RNA-sequencing analysis of differentiated cells. These cells clustered away from undifferentiated iPSCs (Fig. 3a), and they appeared to be largely homogenous with only a few sub-populations not clustered with the main population (Fig. 3b). Blasting the differentiated single cells in a database of human cell atlas containing single cell RNA-sequencing data of various cell types revealed that these iMac cells mainly clustered with macrophages (C0 and C5) or DCs (dendritic cells, C1, C2, C3 and C8) (Fig. 3b, c), with typical genes expressed in each clusters such as CD14 in macrophages and CD1C and HLA-DRB1 in DCs (Supplementary Fig. 4a). Small populations of cells still retained the HSC (hematopoietic stem cell) signature (C9, C7, C4) or differentiated toward other lineages (C6). Trajectory analysis revealed that CAR-iMac cells went through a path from HSC to macrophage and DC cells without major branches (Fig. 3d).

**Fig. 3.**
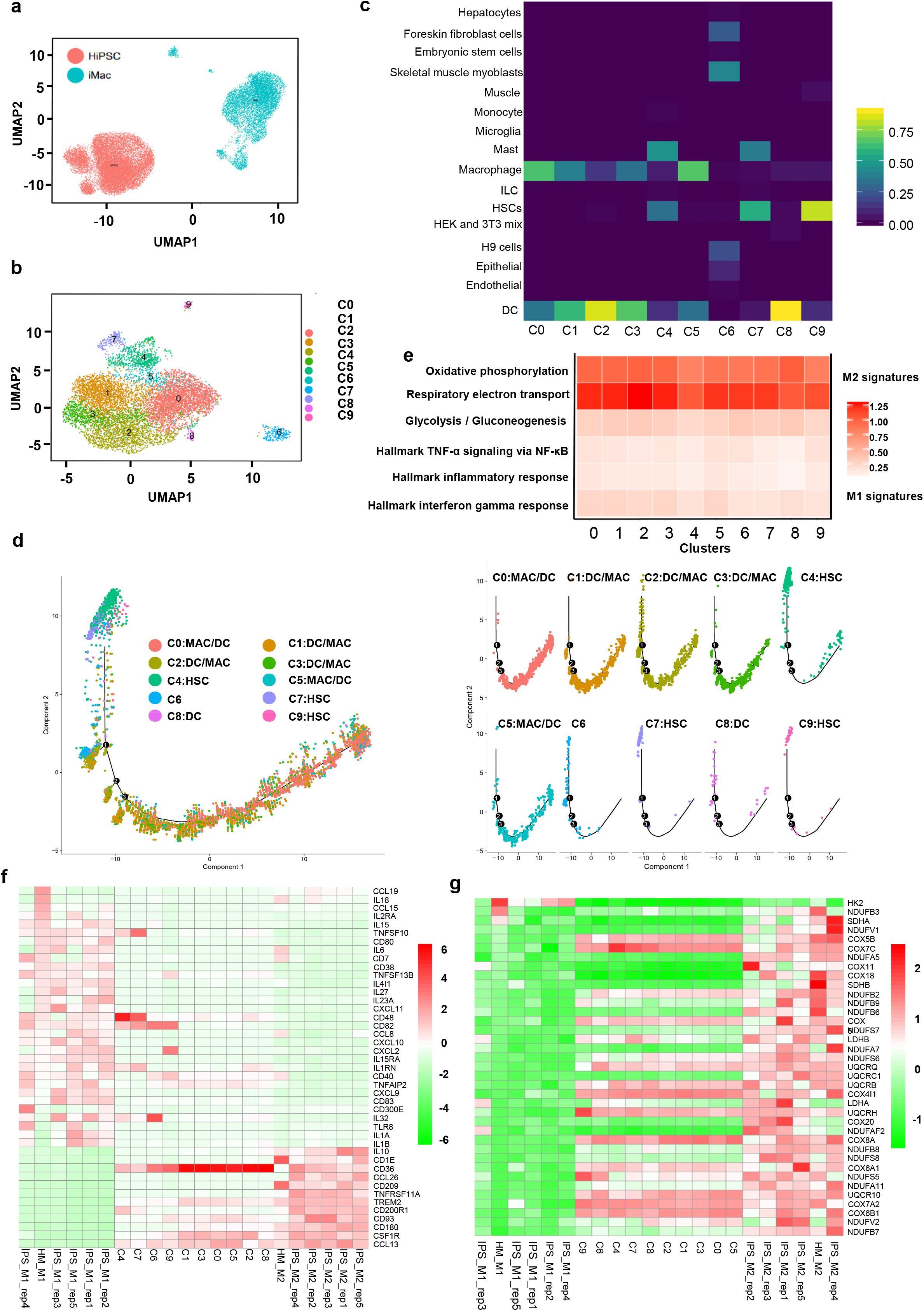
Single cell RNA-sequencing analysis shows CAR-iMac cells are closer to the M2 state on the macrophage polarization spectrum. **a**, UMAP plot showing separation between human iPSCs and CAR-iMac cells. **b**, UMAP plot showing sub-population clustering of CAR-iMac cells. Ten clustered C0-C9 were identified and labeled as 0-9 with different colors. **c**, Heatmap showing blasting the C0-C9 clusters of cells illustrated in **b** against a human single cell atlas database containing single cell RNA-seq data of hundreds of cell types including macrophages (http://scibet.cancer-pku.cn). **d**, Trajectory analysis of differentiated cells along a pseudotime axis. Left panel includes all clusters on the same trajectory map, and right panel is the breakdown of each cluster on the same trajectory. The numbers on the trajectory mean branch points or bifurcating events revealed by the trajectory analysis. MAC/DC means the cluster matches both Macrophage and Dendritic cells. Cluster 6 is not assigned to a specific cell type. **e**, Heatmap showing averaged expression of M1 or M2 signature pathway genes in different clusters of cells illustrated in **b. f, g**, Heatmaps to compare (benchmark) the 10 clusters (C0-C9) of CAR-iMac cells against previously published M1 or M2 polarized macrophages using either top differentially expressed cytokine and chemoattractant genes **f** or using metabolism genes **g** as signature genes. IPS_M1: Human iPS cells differentiated macrophages polarized by IFN-γ and LPS; IPS_M2: Human iPS cells differentiated macrophages polarized by IL-4; HM_M1: Human PBMC-derived macrophages polarized by IFN-γ and LPS; HM_M2: Human PBMC-derived macrophages polarized by IL-4.

In order to determine whether each of these single cells or clusters are more polarized toward the M1 or M2 state, we examined marker gene expression and found high and relatively homogenous expression of M2 genes such as CD206 (Supplementary Fig. 4b). Moreover, all 10 clusters of differentiated cells showed strong signatures of “oxidative phosphorylation” and “respiratory electron transport” which are usually associated with the M2 state, and weak glycolysis or pro-inflammatory signatures associated with the M1 state^16–19^ (Fig. 3e). We further compared cells in the 10 clusters with the LPS/IFN-γ-polarized M1 cells or IL-4/IL-10-polarized M2 cells^14, 15^, by examining differentially expressed cytokine and chemokine genes or M1/M2 associated metabolism genes (Fig. 3f, g), and found that most clusters are more similar to the M2 state, especially when using metabolism genes as markers. Together, these data showed that in the absence of antigen, the first generation of CAR-iMac cells stayed in a state closer to the classically defined M2 along the plastic macrophage spectrum^20–23^, likely because of the presence of M-CSF during the differentiation process^24^.

### CAR-iMac Cells Show Antigen-dependent Functions When Co-cultured with Leukemia and Lymphoma Cells *in Vitro*

In order to test whether iPSC-derived CAR-expressing macrophages assumed the capability to phagocytose tumor cells in an antigen-dependent manner, we incubated the CAR-iMac cells labeled by a green dye with CD19-expressing K562 cells or K562 cells (K562 cell itself does not express CD19) labeled by a red dye. Compared with K562 alone, K562-CD19 cells were more likely to be phagocytosed by CAR-iMac indicated by an increase from 51.9% to 76.4% double positive cells which are the macrophages in the process of phagocytosing tumor cells (Fig. 4a, b). Intracellular signaling such as phosphorylation of ERK and NF-κB P65 proteins were increased in CAR-iMac co-cultured with CD19-expressing K562 cells compared to K562 cells, or to CAR-iMac cells cultured alone (Fig. 4c), indicating antigen engagement promotes activation of macrophage intracellular signaling pathways.

**Fig. 4.**
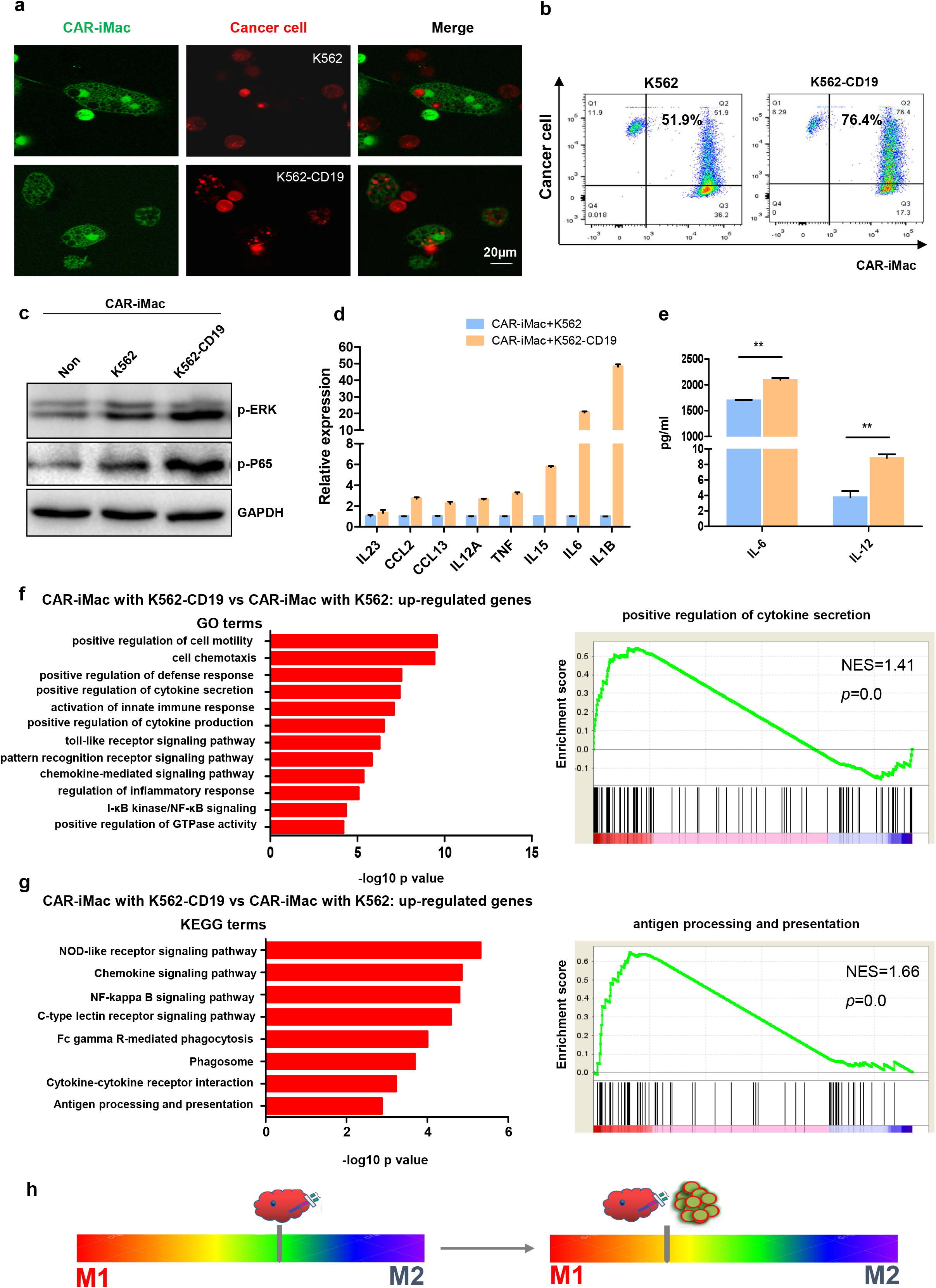
CAR-iMac cells assume antigen- and CAR-dependent macrophage phagocytosis function and pro-inflammatory signatures when co-cultured with liquid cancer cells. **a**, Confocal microscopy pictures showing phagocytosis of K562 or K562-CD19 cells (red) by CAR-iMac cells (green). iMac cells labeled by CMFDA were co-incubated with cancer cells labeled by CTDR for 24 hours before imaging. **b**, Flow cytometry showing phagocytosis of K562 or K562-CD19 cells by CAR-iMac cells. Labeled iMac cells were co-incubated with labeled cancer cells and collected for FACS analysis. Double positive cells represent the iMac cells that engulfed cancer cells. **c**, Western blotting showing phosphorylation of ERK and NF-κB P65 in CAR-iMac cells in the indicated conditions. **d**, qRT-PCR showing cytokine gene mRNA expression when CAR-iMac cells were incubated with K562 or K562-CD19 cancer cells for 24h, cancer cells were washed off before collecting CAR-iMac. n=3, error bar: standard error of the mean. **e**, ELISA assay showing cytokine secretion in the medium when CAR-iMac cells were incubated with K562 or K562-CD19 cancer cells. n=3, error bar: standard error of the mean. **: p < 0.01, one-way ANOVA. **f**, Top GO terms enriched in genes up-regulated in CAR-iMac cells co-cultured with K562-CD19 cells compared with CAR-iMac cells co-cultured with K562 cells without the CD19 antigen. Right panel is GSEA analysis of “positive regulation of cytokine production”. **g**, Top KEGG pathways enriched in genes up-regulated in CAR-iMac cells co-cultured with K562-CD19 cells compared with CAR-iMac cells co-cultured with CD19 cells. Right panel is GSEA analysis of “antigen processing and presentation”. **h**, Schematic diagram showing CAR-iMac cells are more skewed toward M1-like pro-inflammatory state upon encountering the antigen.

One important role of macrophages is to modulate the tumor microenvironment by producing cytokines that either stimulate or suppress T cell activity^9^. In order to harness these features, we first examined cytokine gene expression in CAR-iMac cells, and found antigen-dependent increase of pro-inflammatory cytokine expression, such as *IL1B, IL21, IL12A* and *IL6,* when CAR-iMac cells were incubated with CD19-expressing K562 cells (Fig. 4d), as well as antigen-dependent increase of cytokine secretion in the culture medium such as IL-6 and IL-12 determined by ELISA from the same condition (Fig. 4e). Moreover, transcriptional analysis showed that CAR-iMac cells co-cultured with K562-CD19, in comparison to those with K562, showed strong enrichment of up-regulated genes in GO or KEGG terms of “positive regulation of cytokine secretion”, “antigen processing and presentation” and “Toll-like receptor signaling pathway”, indicating these cells are more wired toward the pro-inflammatory state, when they encounter the antigen (Fig. 4f, g). We also examined another leukemia cell line Nalm6 expressing CD19, and consistently, the CAR-expressing iMac cells showed increased production of pro-inflammatory cytokines and increased level of phosphorylation of ERK, compared to CAR-iMac cells alone (Supplementary Fig. 5a, b), again demonstrating the role of CAR-antigen engagement in conferring the stimulated macrophage activity.

We next investigated the effects of CAR-iMac or iMac cells on a CD19-expressing B cell lymphoma cell line Raji. There is mild enhancement of phagocytosis conferred by the CAR (Supplementary Fig. 5c, d). Moreover, CAR-iMac also had significantly increased pro-inflammatory cytokine gene expression such as *IL18, IL6* and *IL1B,* and other macrophage function-specific genes compared to iMac without a CAR (Supplementary Fig. 5e). When we examined intracellular signaling pathways, we found phosphorylation of AKT, SRC, STAT5 and ERK were all up-regulated when CAR-iMac cells encountered Raji cells (Supplementary Fig. 5f). Taken together, the above data demonstrate that CAR-mediated signaling promotes iMac phagocytosis and tuning toward the pro-inflammatory M1-like state upon encountering the antigen on multiple types of leukemia and lymphoma cells (Fig. 4h).

### CAR-iMac Cells Show Antigen-dependent Functions When Co-cultured with Solid Tumor Cells *in Vitro*

To test iMac cells functions against solid tumors, we replaced the CD19 CAR scFv with a mesothelin scFv in the lentiviral construct, and transduced the new construct to iPSCs followed by differentiating them to CAR (meso)-iMac cells (Fig. 5a). We incubated the CAR (meso)-iMac cells or control iMac cells with mesothelin-expressing OVCAR3 ovarian cancer cells. Compared with control cells, CAR (meso)-iMac showed increased phagocytosis activity against OVCAR3 (Fig. 5b, c). Moreover, transcriptome analysis showed that CAR (meso)-iMac cells also had significantly up-regulated inflammatory and cytokine activity signatures compared to the control when encountering OVCAR3 cells (Fig. 5d, e), which were validated by increased pro-inflammatory M1 cytokine gene expression such as *IL6, IL1A, IL1B,* and *TNF* (Fig. 5f). We also used a mesothelin-expressing pancreatic cancer line ASPC1, and found the same trends of up-regulated phagocytosis, M1 cytokine gene expression, and global inflammatory signatures (Fig. 5g-j). Together, these data demonstrate CAR confers enhanced phagocytosis and pro-inflammatory immune activities when CAR-iMacs are stimulated by antigen-bearing solid tumor cells.

**Fig. 5.**
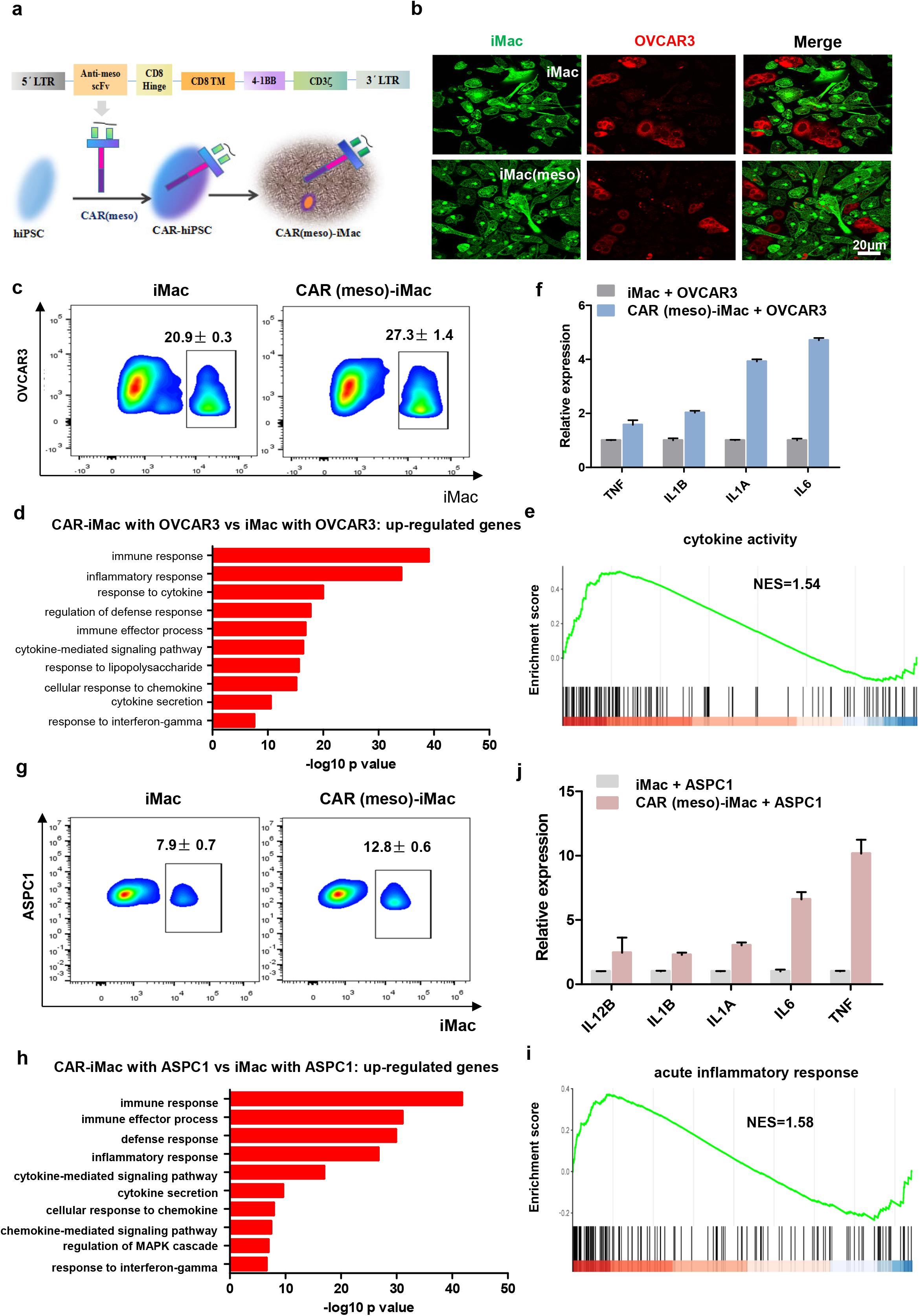
iMac cells assume antigen- and CAR-dependent macrophage phagocytosis functions and pro-inflammatory signatures when co-cultured with solid tumor cells. **a**, A schematic picture to show the lentivirus construct of the CAR against mesothelin CAR (meso), and the procedures to obtain CAR (meso)-iMac cells. **b**, Confocal microscopic images showing phagocytosis of OVCAR3 cells (red) by iMac or CAR (meso)-iMac cells (green). **c**, Flow cytometry showing phagocytosis of OVCAR3 ovarian cancer cells by iMac or CAR (meso)-iMac cells. Labeled iMac cells were co-incubated with labeled cancer cells and collected for FACS analysis. Squared parts represent the cancer cells engulfed by iMac cells. n=3. **d**, GO term analysis with RNA-seq data showing the up-regulated genes in CAR (meso)-iMac cells compared with iMac cell when both cells were co-cultured with OVCAR3 cancer cells. **e**, GSEA analysis of representative cytokine activity gene set showing it is up-regulated in CAR (meso)-iMac cells compared with iMac cell when both cells were co-cultured with OVCAR3 cells. NES: normalized enrichment score. **f**, qRT-PCR showing cytokine gene mRNA expression when iMac or CAR (meso)-iMac cells were incubated with OVCAR3 cells for 24h. Cancer cells were washed off before collecting iMac or CAR-iMac cells. n=3. Error bar: standard error of the mean. **g**, Flow cytometry showing phagocytosis of ASPC1 pancreatic cancer cells by iMac or CAR (meso)-iMac cells. n=3. **h**, GO term analysis showing the up-regulated genes in CAR (meso)-iMac cells compared with iMac cell when both cells were cocultured with ASPC1 cells. **i**, GSEA analysis of representative acute inflammatory response gene set showing it is up-regulated in CAR (meso)-iMac cells compared with iMac cell when both cells were co-cultured with ASPC1 cells. **j**, qRT-PCR showing cytokine gene mRNA expression when iMac or CAR (meso)-iMac cells were incubated with ASPC1 cells for 24h. Cancer cells were washed off before collecting CAR-iMac. n=3, error bar: standard error of the mean.

### CAR-iMac Cells Show in *Vivo* Phagocytosis and Anti-Cancer Cell Function

In order to detect phagocytosis functions of CAR-iMac cells *in vivo,* we injected GFP-expressing Raji cancer cells and CAR-iMac cells to NSG mice intravenously, and after three days, the sorted human CD11b^+^ (from CAR-iMac cells) and GFP^+^ (from cancer cells) double positive cells were examined with Giemsa staining (Fig. 6a). The macrophages showed multiple nuclei, suggesting *in vivo* phagocytosis took place in the injected mice (Fig. 6a). To evaluate the anti-cancer cell function of these CAR-iMac cells *in vivo*, we intraperitoneally injected leukemia cells expressing a luciferase gene into NSG mice, and after tumors formed, mice bearing comparable tumor loads were separated into two groups for treatment and controls. In order to achieve better anti-cancer cell outcome, we further polarized the CAR-iMac cells toward M1 by treating them with IFN-γ for 24 hours (Supplementary Fig. 6a), and then one group of mice was injected intraperitoneally with the polarized CAR-iMac cells for treatment (Fig. 6b). The treated mice showed smaller tumor size on day 10, with two mice having complete remission (Fig. 6c). For solid tumors, we inoculated a high mesothelin-expressing ovarian cancer cell line HO8910 intraperitoneally to NSG mice, followed by intraperitoneally injecting IFN-γ-treated CAR (meso)-iMac cells (Fig. 6b). Two out of the five treated mice showed tumor remission (Fig. 6d), and four out of five mice in the treated group lived for three weeks but only two mice in the control group (Fig. 6d). The treatment did not work to the same extent in all mice, likely because not all the plastic iMac cells preserved in the same polarized state in the complex and sometime immunosuppressive *in vivo* microenvironment in all mice. Nevertheless, these data demonstrate CAR-iMac cells showed some anti-cancer cell functions *in vivo*, but their efficiency needs to be further improved by designing more effective CARs or further modifying the CAR-iMac cells to stay constitutively in M1 state.

**Fig. 6.**
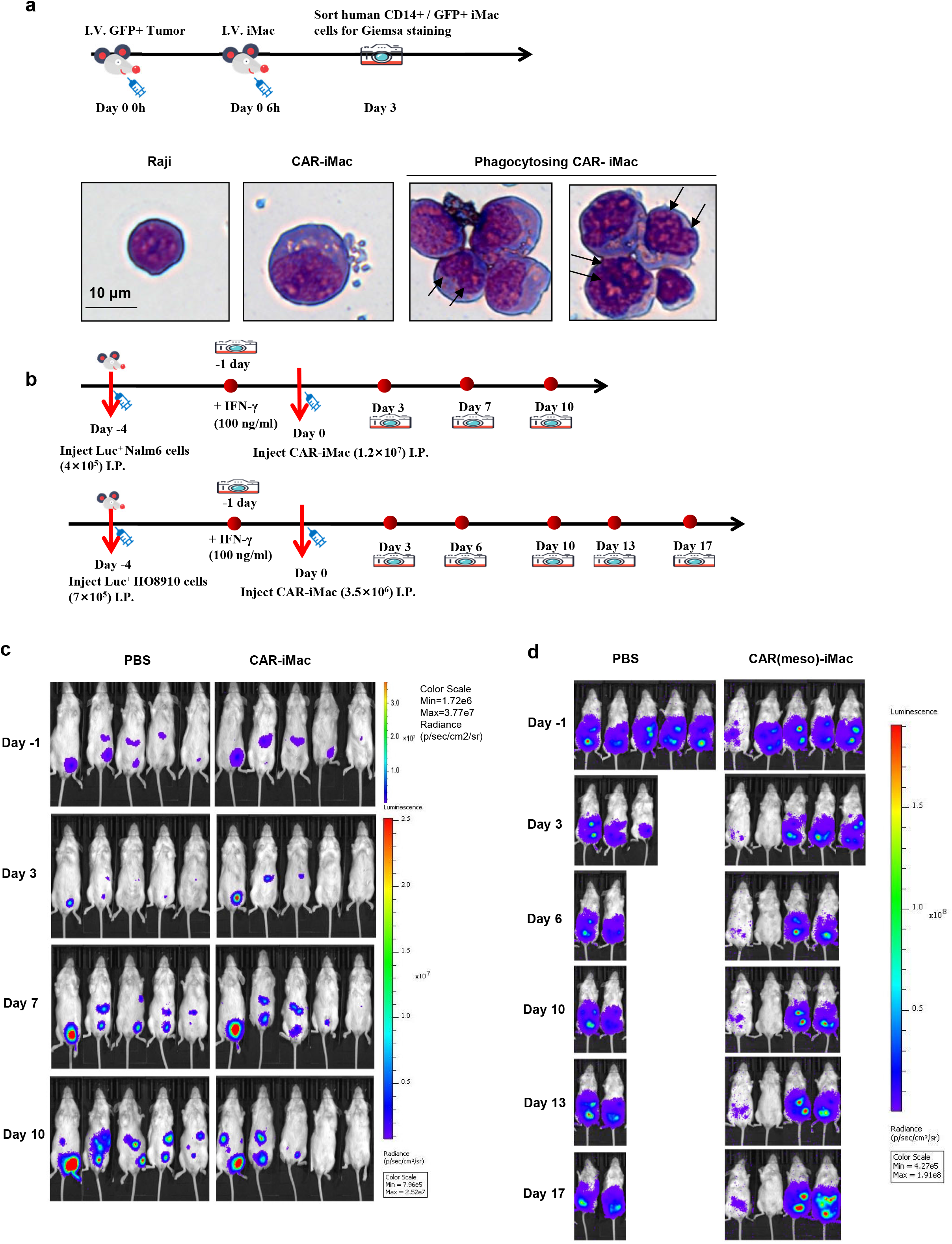
CAR-iMac cells showed *in vivo* phagocytosis function and anti-tumor effect. **a**, Giemsa staining of FACS-sorted human CD11b and GFP double positive cells showing that CAR-iMac cells phagocytosed Raji cells *in vivo.* 4×10^5^ GFP-labeled cancer cells were injected intravenously, and 6 hours later, 1×10^6^ CAR-iMac were injected intravenously. At day 3, cells were sorted for staining. Black arrows point to the multiple nucleus in the isolated macrophages. **b**, A schematic picture of *in vivo* anti-tumor studies using luciferase (luc)-labeled CD19-expressing Nalm6 leukemia cells or mesothelin-expressing HO8910 ovarian cancer cells in NSG mice treated with CAR-iMac cells. **c**, Bioluminescent imaging pictures showing tumor development on the indicated days in mice inoculated with Nalm6 leukemia cells and treated with iMac cells or PBS as described above. **d**, Bioluminescent imaging pictures showing tumor development on the indicated days in mice inoculated with HO8910 ovarian cancer cells and treated with iMac cells or PBS.

## Discussion

Recently, iPSC-differentiated CAR-expressing T cells and NK cells have been reported to have potent cytotoxic activity against cancer cells, and they represent a new family of engineered stem cell-derived immune cells for CAR therapies^5, 6^ A new member of this family, CAR-iMac has three advantages. First, CAR-iMac can be utilized to specifically “eat” tumor cells or to alter the tumor microenvironment, especially when its function is antigen-dependent, providing a tool to directly kill tumor cells or to modulate a specific niche at the interface of tumor and immune cells. Second, increasingly complicated genome engineering approaches can be applied to iPSCs, followed by single clone expansion and thorough characterization to obtain a clonal and genetically homogenous population with minimal off-target risk. Third, the seed iPSC clone can be expanded in large scale, and when combined with the recently reported techniques in making immune-tolerant universal iPSCs^25^–^27^, CAR-iMac can be developed into an off-the-shelf product with unlimited sources. The last two points are particularly useful when the effector cells from primary cell sources are difficult to engineer and to expand which is the case for PBMC-derived macrophage cells. Thus, CAR-iMac is an excellent platform for engineering-friendly and expandable macrophage cells, and a valuable addition to other iPSC differentiated immune cells for further cancer immunotherapies.

There are several general challenges of using iPSC differentiated cells for clinical applications, such as undifferentiated cell can form teratomas, yield of differentiation and heterogeneity of differentiated cells. We have achieved high differentiation efficiency with a yield of ∼10-100 myeloid cells per starting iPSC. Even though certain cells remained some gene expression signature of stem/precursor cells, we have not seen teratomas formation when we transplanted the bulk differentiated cells into mice. In terms of heterogeneity, differentiated cells collected from the floating shedded cells are primarily positive for macrophage markers such as CD14 and CD11b (Fig. 1c). More careful examination with single cell RNA-sequencing data revealed a clear trajectory of differentiation toward macrophages and dendritic cells (Fig. 3c), which by themselves share overlapping transcriptome signatures and functions, and can be named altogether as “mononuclear phagocyte system (MPS)” cells^28^. Notably, in CD19 CAR-T cell therapies or immune checkpoint therapies, combinations of CD8^+^ and CD4^+^ T cells have superior anti-cancer cell activity^29–31^. It is thus conceivable that MPS cells may achieve synergistic effects from different subsets of cells by simultaneously phagocytosing tumor cells or presenting antigens to T cells. Therefore, it will be important to compare the purified versus mixed myeloid products for their efficacy, persistence and toxicity in future studies. It will also be of high interest to use humanized mouse models to study the role of differentiated DC-like cells in tumor antigen presentation and T cell stimulation.

Another challenge to solve in order to take our findings to the clinic is to balance the anti-cancer cell functions from pro-inflammatory cytokines and the toxicity induced by the same cytokines such as IL-1 and IL-6^32, 33^. In our system, we achieved antigen-dependent induction of these cytokines, so that they can only be released in large amount within a restricted niche of antigen exposure. Indeed, our first generation of iMac cells were in a state closer to the immune-suppressive M2 state and were tuned toward M1 after exposing to antigens, which suggests that the cells may travel safely in the blood vessel, and may not become inflammatory until it reaches to the tumor microenvironment (Fig. 4h). Another potential solution is to engineer molecular switches to enable CAR or cytokine expression under control of endogenous promoters^34^or exogenously administered small molecules^35^ so that inflammatory cytokine expression is controlled by the cell state or controlled in a temporal way. Moreover, further engineering the CAR intracellular domain to confer phagocytosis function without eliciting inflammation-stimulating cytokines, such as using intracellular domain of FcγR^36^, or knocking out endogenous inflammation-stimulating cytokine genes, are all possible solutions to explore. Besides the safety issues, we explored the anti-tumor effect in animal models of both liquid and solid tumors. The data showed that our first generation of CAR-iMac had some effect. While our paper is being reviewed, Klichinsky *et. al* reported PBMC-derived macrophage transduced with a CAR through adenovirus showed strong *in vivo* anti-tumor effect, and one key reason is the adenoviral vector Ad5f35 used imparted a sustained pro-inflammatory M1 phenotype^37^. Thus, there are many directions to improve the efficacy of our first generation of CAR-iMac along this line. Our current iMac differentiation protocol contains M-CSF which is not only required to support myeloid cell survival, but also unavoidably biases the differentiation toward M2 state. Even though we added IFN-γ to polarize it to M1 state and CAR signaling tuned it toward M1, but the epigenetic memory inherited may influence the ultimate state, particularly for the plastic macrophage in the complicated *in vivo* microenvironment. Thus, a constitutive M1 signaling is required to guarantee the cells to be constantly pro-inflammatory and anti-tumor. With the iMac system, this can be achieved by inserting certain M1 cytokine genes in the genome of iPSC seed cells, or by further optimizing the intracellular domain of CAR to confer M1 signaling, similar as the second signal provided by the “co-stimulatory” domain from CD28 and 4-IBB in the second generation of CAR-T cells^38, 39^.

Together, our data provided a proof-of-principle platform of iPSC-based, engineering-friendly macrophage for adoptive transfer immune cell therapy, and suggest that further development and optimization of CAR-iMac is warranted. Its kaleidoscopic applications in various contexts of liquid and solid tumors, and in combination of genome engineering or in combination with other types of immunotherapies are full of great promises for further immune cell therapies.

